# Dopamine mediates the bidirectional update of interval timing

**DOI:** 10.1101/2021.11.02.466803

**Authors:** Anthony M.V. Jakob, John G. Mikhael, Allison E. Hamilos, John A. Assad, Samuel J. Gershman

## Abstract

The role of dopamine as a reward prediction error signal in reinforcement learning tasks has been well-established over the past decades. Recent work has shown that the reward prediction error interpretation can also account for the effects of dopamine on interval timing by controlling the speed of subjective time. According to this theory, the timing of the dopamine signal relative to reward delivery dictates whether subjective time speeds up or slows down: Early DA signals speed up subjective time and late signals slow it down. To test this bidirectional prediction, we reanalyzed measurements of dopaminergic neurons in the substantia nigra pars compacta of mice performing a self-timed movement task. Using the slope of ramping dopamine activity as a read-out of subjective time speed, we found that trial-by-trial changes in the slope could be predicted from the timing of dopamine activity on the previous trial. This result provides a key piece of evidence supporting a unified computational theory of reinforcement learning and interval timing.

## Introduction

How does dopamine (DA) influence time perception? This question has been an active subject of debate. While some researchers have found that DA increases the rate at which subjective time progresses (Lake and Meck, 2013; Maricq et al., 1981; Maricq and Church, 1983), others have found the exact opposite effect (Soares et al., 2016). Recent work has developed a coherent framework to explain these phenomena (Mikhael and Gershman, 2019), which relates these timing effects to the role of DA in signaling reward prediction error (RPE; for reviews, see Gershman et al., 2014; Petter et al., 2018).

According to the RPE hypothesis, DA reports the difference between received and expected reward. In a seminal experiment, Schultz et al. (1997) presented monkeys with repeated rewards (after a fixed delay from a cue) and simultaneously recorded from putative DA neurons in the midbrain. The authors found that an unexpected reward elicited a burst of DA neuron activity, but that, when the reward was expected, it no longer elicited DA neuron activity. Furthermore, a reward omission at the time of expected reward elicited a *dip* in activity. These experimental observations are consistent with the RPE hypothesis, and have been buttressed by several decades of research (e.g., Eshel et al., 2015; Glimcher, 2011; Niv and Schoenbaum, 2008; Schultz et al., 1997; Steinberg et al., 2013). The computational importance of this hypothesis is due to the role of RPE in reinforcement learning (RL) algorithms, specifically the temporal difference learning algorithm (Sutton, 1988; Sutton and Barto, 2018). An agent can use RPEs to learn long-term reward predictions: Unexpected rewards indicate that the agent should increase its future expectation of reward, while omissions of expected rewards indicate that the animal should decrease its future expectation of reward.

The RPE hypothesis does not by itself explain the role of DA in interval timing, since it is compatible with many different assumptions about the representation of time (Starkweather et al., 2017; Daw et al., 2006; Ludvig et al., 2008). However, the choice of time representation can have a dramatic influence on the effectiveness of RL algorithms. If there is some limit on the precision with which time can be represented, then the limited representational capacity should be concentrated on time scales (or more generally time intervals) that are important for reward prediction. Since animals need to deal with multiple time scales for different tasks, this representation should be rescalable. For example, if time is represented by the firing rate of “time cells” tuned to particular time intervals (e.g., MacDonald et al., 2011; Salz et al., 2016; Tiganj et al., 2017; Bright et al., 2020), then the tuning functions should stretch or compress if the task-relevant interval is increased or decreased, respectively. Evidence for task-dependent rescaling has been reported in both striatum (Mello et al., 2015) and hippocampus (Shimbo et al., 2021).

Mikhael and Gershman (2019) formalized this rescaling idea in a temporal difference learning model of DA. The key idea was to treat the time scale of the temporal representation as a parameter that could be adjusted by the RPE signal. In this way, DA could modify the speed of subjective time in order to optimize reward prediction. In particular, the model predicted a bidirectional plasticity rule for the timing parameter: Positive RPEs that occur *before* expected reward delivery should tend to *increase* the speed of subjective time, while positive RPEs that occur *after* expected reward delivery should *decrease* the speed of subjective time. Mikhael and Gershman (2019) showed that this model could account for a number of dopaminergic effects on interval timing behavior.

In this paper, we undertake a more direct test of the bidirectional plasticity hypothesis, using DA measurements collected from mice performing a self-timed movement task (Hamilos et al., 2020). In this task, mice received a reward for licks performed after a fixed interval. Even after extensive training, the authors observed ramping DA signals and variable trial-to-trial lick times. Furthermore, the authors found that steeply rising DA ramps preceded early lick times and slowly rising DA ramps preceded late lick times. Based on our earlier theoretical work (Gershman, 2014; Kim et al., 2020; Mikhael et al., 2021), we argue that the slope of DA ramps is a proxy for the speed of subjective time. We then ask whether the timing of DA activity relative to the time of reward delivery predicts the ramp slope on the subsequent trial in accordance with the bidirectional plasticity rule.

## Methods

### The computational problem

We construe animals as facing the problem of learning to predict long-term reward, or *value*, defined as the expected discounted future return (cumulative reward):

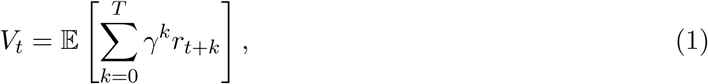

where *t* indexes intra-trial time (*t* = 0 corresponds to trial onset), *r*_*t*_ is the reward received at time *t, T* is the trial duration, and *γ* ∈ (0, 1) is a discount factor. In Hamilos et al. (2020), the animal receives a single reward *r* at time *T* in each trial, so Equation (1) can be simply written as:

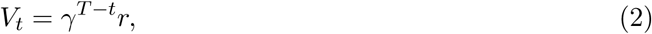

The value function and RPE are illustrated in Figure 1A. If, as commonly assumed, the rewards follow a Markov process, then Equation (1) can be written recursively:

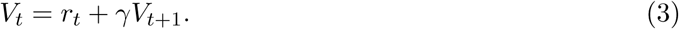

**Figure 1:**
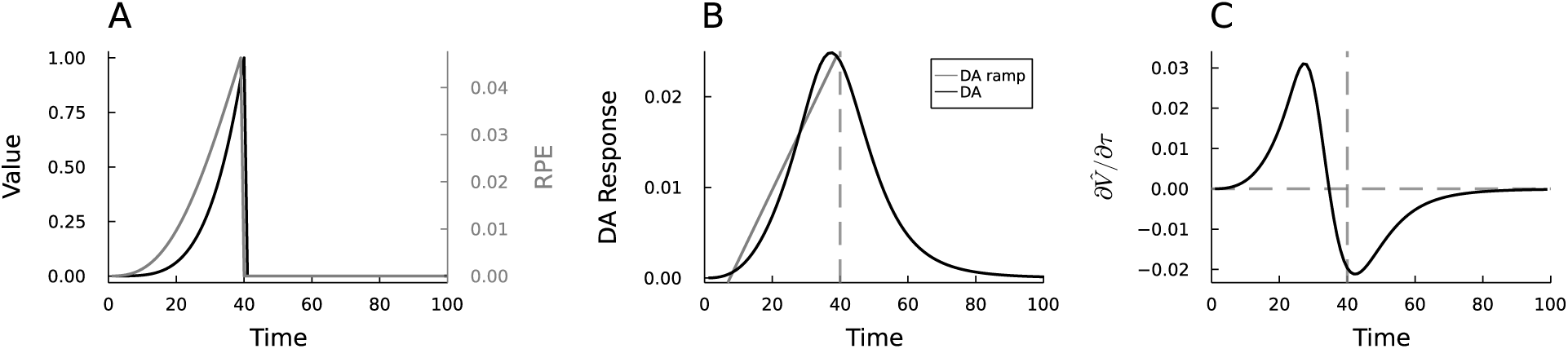
Simulations of ramping RPEs with a temporal difference learning model. (A) Convex value function (black) and ramping RPE (gray). (B) Simulated DA signal (black) and estimated DA ramp (linear regression between trial start and reward delivery, gray). The DA signal corresponds to the RPE under temporal uncertainty (Methods). (C) Partial derivative of the estimated value function with respect to time, which gives the bidirectional update rule of the pacemaker rate *η* its qualitative shape.

This recursive expression is known as the Bellman equation (Bellman, 1957), and is the basis for efficient RL algorithms such as temporal difference learning (Sutton, 1988).

### Temporal difference learning model

To learn the value function *V*_*t*_, we first define a parametric function class and then present a learning algorithm that adjusts the parameters to minimize the discrepancy between the estimator and the true value function. A standard parametrization is the linear function approximator, which approximates the value function as a linear projection of time-varying features (Schultz et al., 1997; Ludvig et al., 2008, 2012):

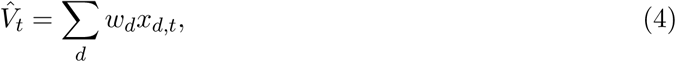

where *x*_*d,t*_ is the *d*^*th*^ feature at time *t*, and *w*_*d*_ is the feature weight. For example, a feature may represent the presence (*x*_*d,t*_ = 1) or absence (*x*_*d,t*_ = 0) of a stimulus at time *t*. Alternatively, it may represent the physical proximity to a reward location.

The weights *w*_*d*_ are updated by gradient descent to reduce the mismatch between *V*_*t*_ and 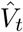:

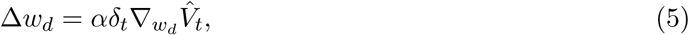

where *α* ∈ (0, 1) is the learning rate, 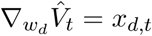 is the gradient of 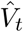 with respect to the weight *w*_*d*_, and *δ*_*t*_ is the reward prediction error (RPE):

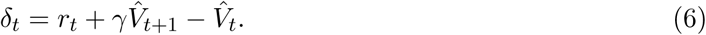

Notice that *δ*_*t*_ equals the mismatch between the agent’s estimates of the right-hand side and left-hand side of Equation (3). When *δ*_*t*_ = 0 on average, 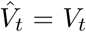, and hence the value is well-learned. Otherwise, the agent continues to update 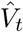 to minimize *δ*_*t*_.

The shape of *δ*_*t*_ after a task is well-learned will depend on the choice of features. For instance, Gershman (2014) showed that, for a single feature taking sufficiently convex shape across states, *δ*_*t*_ will exhibit the shape of a ramp (see also Morita and Kato, 2014; Lloyd and Dayan, 2015; Mikhael et al., 2021, for alternative approximation architectures that result in ramps, such as time cells). For simplicity, we will assume in what follows a single feature *x* encoding a subjective representation of elapsed time since the beginning of the trial. Subjective time is assumed to be a convex function of objective time, which will produce ramping (Figure 1A; a mathematical analysis of this point appears below).

It is important to note that perfectly learning a value function depends on having a perfect internal clock (i.e., subjective and objective time coincide). Instead, animals are noisy timers, and are furthermore subject to Weber’s law, which asserts that the standard deviation of an animal’s temporal estimate increases linearly with the elapsed time (Church and Meck, 2003; Gibbon, 1977; Staddon, 1965). This has the effect of ‘blurring’ the value function in proportion to the animal’s temporal uncertainty. Because the RPE is a function of value, it too gets blurred, and this blurring determines the shape of the ramp (Figure 1B). Specifically, the predicted DA response is computed as the convolution of the RPE with a Gaussian temporal uncertainty kernel determined by Weber’s law:

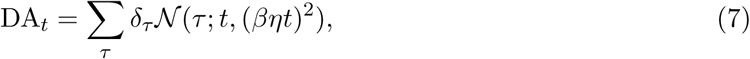

where *β* is the Weber fraction.^1^ In our previous work (Mikhael et al., 2021), we showed that temporal uncertainty can explain diverse DA dynamics across different tasks, including positive ramps, negative ramps, flat functions, and even non-monotonic functions.

The key addition of the model presented in Mikhael and Gershman (2019) is to account for the role of DA in modulating the speed of subjective time. We formalize this speed variable as a parameter *η* that rescales the relationship between objective and subjective time: *τ* = *ηt*. Thus, when *η* increases, subjective time (*τ*) runs faster. Importantly, we can view *η* as another parameter in the function approximation architecture, and optimize it via gradient descent just as we did for the weights:

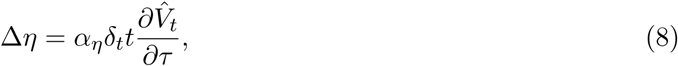

where *α*_*η*_ is the learning rate. Note here that the derivative of 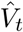 is greater than zero roughly before reward delivery but less than zero afterwards. It follows that the contribution of the RPE is bidirectional: DA signals occurring before reward time should increase *η*, and DA signals occurring after reward time should decrease it (Figure 1C).

### Choice of feature shape

Our choice of feature *x* results in a ramping RPE. To see this, note that an RPE ramps if and only if 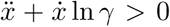 (Mikhael et al., 2021). Intuitively, by Equation (6), *r*_*t*_ = 0 during the trial but prior to receiving reward. With a single feature, it follows that 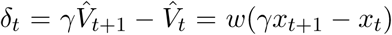. Because *γ* is close to 1, the term in the parentheses is approximately the derivative of *x*. This term, and hence the RPE, ramps when its own derivative is positive, i.e., when the second derivative of *x* is positive 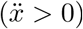. The second term in our exact requirement accounts for the more general case when *γ* is not equal to 1 (see Mikhael et al., 2021, for a full derivation of this result). Using our choices of *x* and *γ* (specified below), the requirement is satisfied for *t <* 58, which is a superset of the temporal domain chosen for our simulations.

### Simulation parameters

We have chosen *γ* = 0.95, *T* = 40, *β* = 0.2, *r*_*T*_ = 1 at time *t* = *T* and *r*_*t*_ = 0 otherwise, *τ* = *t, α*_*η*_ = 0.01, and a single feature *x*_*t*_ = *kt*^4^ if *t* ≤ *T*, and 0 otherwise, with *k* = *r*_*T*_ *T* ^*−*4^.

### Data analysis

We computed the DA dF/F signal from the raw GCaMP6f measurements according to Hamilos et al. (2020). We then divided each trial into *n* time-bins. We chose *M* = 20 time-bins of length 0.85s each to account for the trial length of 17s. We aligned the time-bins around the first-lick time in each trial *n* and computed the average DA level *D*_*n,m*_ within each time-bin *m*. We computed the baseline DA level for each trial, defined as the average DA level between lamp-off (a signal indicating the imminence of the cue, see Hamilos et al. (2020)) and cue, and subtracted it from each corresponding time-bin. Then, we computed the DA ramp slope *s*_*n*_ during the trial by fitting a straight line to the DA signal from 0.7s post-cue to 0.6s pre-lick. These buffer lengths are taken from Hamilos et al. (2020) to eliminate the effect of motion-induced transients in the signal. We then defined *a*_*n*_ = *s*_*n*+1_ − *s*_*n*_, the difference in DA slope between the current trial *n* and the next trial *n* + 1, which is a neural proxy for the change in *η* from the current trial to the next. We then solved the linear system *Db* = *a*, where *b* is the contribution of each bin to the change in DA ramp slope. The solution to this optimization problem (equivalent to maximum likelihood estimation of a linear regression model) is 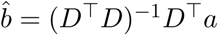. This analysis was done for each mouse individually as well as on pooled data.

### Source code

All simulations and analyses were performed using Julia, version 1.6.2. Source code can be found at https://github.com/antjak/dopa-rpe-interval-timing.

## Results

Hamilos et al. (2020) trained mice to perform an interval timing task by initiating a self-timed lick at least 3.3 seconds after a start-timing cue. Despite highly variable lick times from trial to trial, the authors found that DA signals ramped up during the self-timed interval following the start-timing cue. Crucially, they found that the DA ramp slope was highly predictive of lick time, with larger slopes being associated with earlier lick times. They also found that higher baseline DA levels correlated with greater ramp slopes and earlier lick times, consistent with the view that higher DA levels lead to faster clocks.

To examine our prediction of a bidirectional effect of DA on the speed of subjective time, we reanalyzed the data from Hamilos et al. (2020). Using the linear regression model detailed in the Methods, we studied the association between DA levels at particular points in time during a trial and the ramp slope (a measurable proxy for the speed of subjective time) on the subsequent trial. In this way, we could extract a detailed temporal plasticity function and compare it to the theoretical plasticity function (Figure 1C).

Figure 2A shows the estimated regression coefficients for each time-bin. Consistent with our model predictions, the estimated coefficients revealed that early DA signals in a trial had a positive effect on the change in ramp slope, and late signals had a negative effect. In other words, an increase in DA activity shortly after cue presentation resulted in an increase in ramp slope on the next trial, whereas an increase in DA activity shortly after licking resulted in a decrease in ramp slope on the next trial.

**Figure 2:**
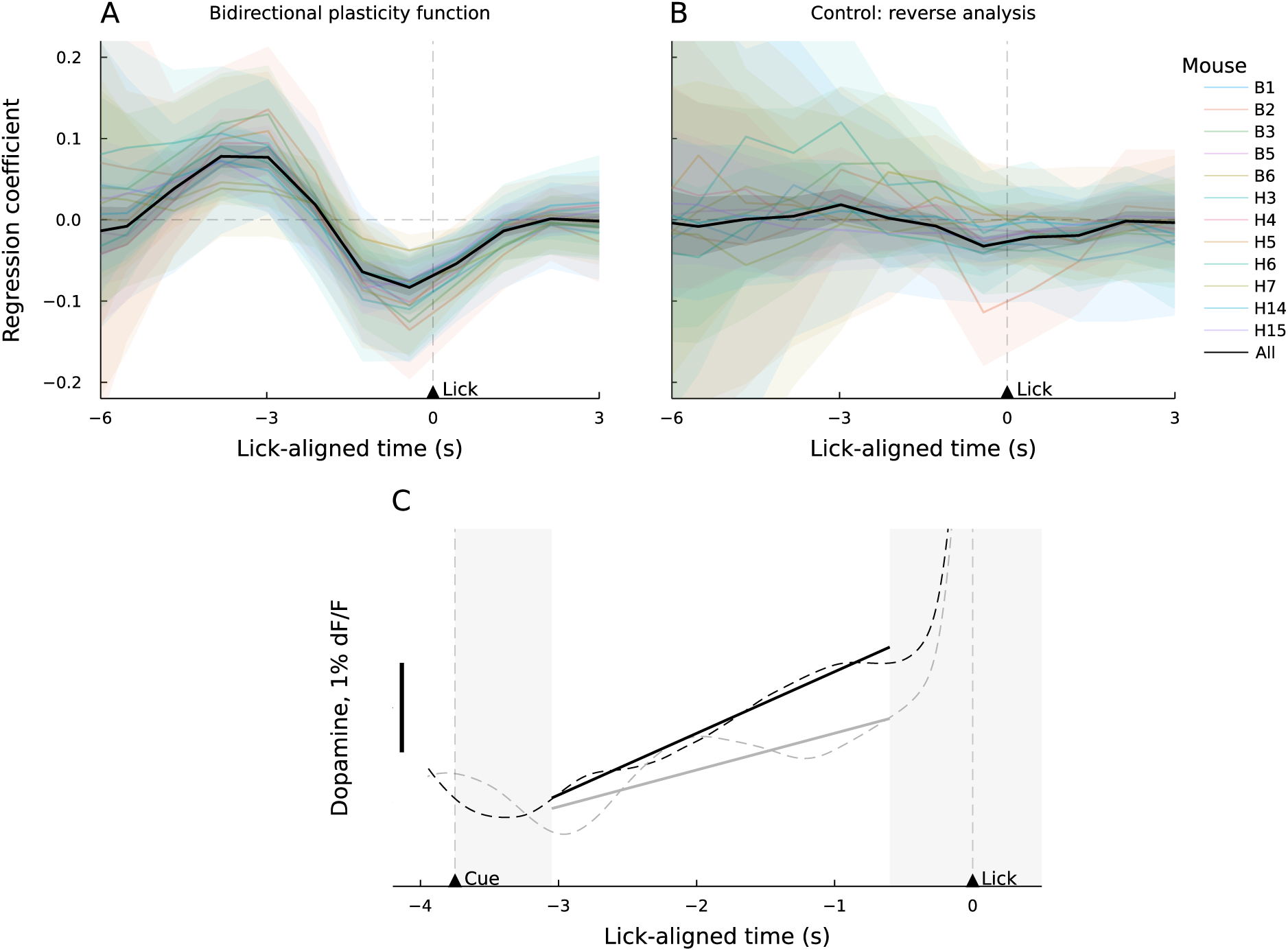
Bidirectional update rule. (A) Empirical bidirectional plasticity function for rewarded trials with a lick 3.3-7s post-cue, for each mouse (colors) and for pooled data (black), smoothed with a 1.7s moving average filter. The function’s qualitative shape is not sensitive to the precise choice of lick interval. Shaded area corresponds to standard error. Regression coefficient of bin after cue and bin immediately before lick are statistically different (*t*(11) = 8.7, *p <* 10^−5^). (B) Empirical plasticity function for reversed trial order to rule out a possible slow temporal confound, smoothed with a 1.7s moving average filter. Regression coefficient of bin after cue and bin immediately before lick are not statistically different (*t*(11) = −0.4, *p* = 0.68). (C) Average DA signal for trials with a lick 3.5-4s post-cue, smoothed with a 400ms moving average kernel and classified by DA level on the previous trial—high DA around the cue (black) and high DA around licking (gray). The dashed lines correspond to the average DA signal, while the thick lines are fitted to the signal between the gray rectangles, which represent the buffers after the cue and before the lick, as given in Hamilos et al. (2020). A high-DA-around-cue condition (*n* = 1882) corresponds to a steeper DA ramp slope on the next trial, as compared to a high-DA-around-lick condition (*n* = 3668; *t*(5548) = 7.2, *p <* 10^−12^).

Due to the slow drift in the behavioral timing distribution occurring between the beginning and end of sessions, higher baseline amplitude at the beginning of the session may lead to steeper slopes on nearby trials generally, without any causal effect. Although baseline normalization of activity on each trial should diminish the effect of slow drift, it is possible that residual drift is driving our results. We reasoned that if the slow drift hypothesis is correct, then it should also produce the same results when run on trials in the reverse order. We therefore reran the regression analysis on the reversed sequence of trials, which eliminated the relationship between within-trial DA signaling and ramp slope change (Figure 2B). This analysis, coupled with baseline normalization, rules out the slow temporal confound.

Figure 2C illustrates how ramp slope changes as a function of DA activity at different points during the previous trial. When DA activity is high following cue presentation, the ramp on the next trial tends to be steeper compared to when DA activity is high immediately before licking. Our model asserts that this difference arises from the proposed bidirectional plasticity rule.

## Discussion

By reanalyzing recordings of dopaminergic neurons in mice performing a self-timed movement task (Hamilos et al., 2020), we have shown that DA has a bidirectional effect on the speed of subjective time. We showed that the contribution of DA on the current trial to the change in DA ramp slope (a proxy for the speed of subjective time) on the next trial exhibits the predicted bidirectional shape: DA signals occurring before reward time tend to increase the DA ramp slope on the next trial, and those occurring after reward time tend to decrease it, consistent with our RL theory of temporal optimization (Mikhael and Gershman, 2019). This theory was previously invoked by Hamilos and Assad (2020) to explain variability in ramp slope due to time rescaling, but that study left open the question of why time rescaling itself should vary across trials. Here we address this question by showing how time rescaling can be endogenized by a model that optimizes the rescaling parameter using temporal difference learning.

For simplicity, we have chosen a feature in our temporal difference model that produces ramps. However, the cause of ramps—and how they relate mechanistically to the flow of time—remains an open question. Indeed, DA ramps have been observed in various operant conditioning tasks, both during the pre-action period (Totah et al., 2013) as well as during action execution (Howe et al., 2013). Ramps have furthermore been observed in classical conditioning tasks that provided cues indicating proximity to reward (Kim et al., 2020). Recent work has suggested that these ramps occur as a consequence of sensory feedback (Mikhael et al., 2021), although they may also be captured by a “forgetting” mechanism within an RL framework (i.e., a decay term in the value update; Morita and Kato, 2014), or by state-dependent biases such as an overestimation of time or distance to reward, if the biases decrease with proximity to the reward (Mikhael et al., 2021).

Our model of timing optimization by RL can potentially be related to several existing models of interval timing. In the striatal beat frequency model, cortical neurons are assumed to fire in an oscillating pattern with different phases (Matell and Meck, 2004). It follows that the neurons active during both the reward-predicting cue and the reward represent a neural code for the interval to be timed. Assuming that DA affects the firing frequency of the cortical oscillators, our bidirectional update rule provides a compatible extension to this model to account for interval timing modulation effects.

Alternatively, in pacemaker-accumulator models, time is represented by counting the number of ticks emitted by a noisy clock (Gibbon et al., 1997; Zakay and Block, 1997). Given the similarity between the ticking of the clock and the successive transition from state to state—typical for a RL model—as a representation of the passing of time, our model provides a natural extension to the PA framework: By letting the rescaling parameter *η* influence the speed of the clock or the tick number threshold, DA-mediated interval timing modulation can be accounted for. Despite the differences between both classes of models of timing presented here, it is interesting to note that the parametrized rescaling of a quantity through a bidirectional plasticity rule will endow the model with the ability to accurately account for interval timing modulation effects.

In conclusion, we have shown here that RL and interval timing are critically linked by a common dopaminergic mechanism. To our knowledge, this is the first theory that captures the bidirectional effect of DA on interval timing. More broadly, the idea that prediction errors can drive representation learning may extend beyond interval timing to other domains (Alexander and Gershman, 2021). An important project for future work will be to examine empirically whether the same dopaminergic signal serves this function across domains.

## Footnotes

1 The assumption of Weber noise is not necessary for the results we present in this paper, but we include it here for consistency with past work.

